# Seascape genomics reveals candidate molecular targets of heat stress adaptation in three coral species

**DOI:** 10.1101/2020.05.12.090050

**Authors:** Oliver Selmoni, Gaël Lecellier, Hélène Magalon, Laurent Vigliola, Francesca Benzoni, Christophe Peignon, Stéphane Joost, Véronique Berteaux-Lecellier

**Affiliations:** Laboratory of geographic information systems, Ecole Polytechnique Federale de Lausanne, Lausanne Switzerland; UMR250/9220 ENTROPIE IRD-CNRS-Ifremer-UNC-UR, Labex CORAIL, Nouméa, New Caledonia; UVSQ, Université de Paris-Saclay, Versailles, France; UMR250/9220 ENTROPIE IRD-CNRS-Ifremer-UNC-UR, Labex CORAIL, Université de La Réunion, St Denis, France; Red Sea Research Center, Division of Biological and Environmental Science and Engineering, King Abdullah University of Science and Technology, Thuwal, Saudi Arabia

## Abstract

Anomalous heat waves are causing a major decline of hard corals around the world and threatening the persistence of coral reefs. There are, however, reefs that had been exposed to recurrent thermal stress over the years and whose corals appeared tolerant against heat. One of the mechanisms that could explain this phenomenon is local adaptation, but the underlying molecular mechanisms are poorly known.

In this work, we applied a seascape genomics approach to study heat stress adaptation in three coral species of New Caledonia (southwestern Pacific) and to uncover molecular actors potentially involved. We used remote sensing data to characterize the environmental trends across the reef system, and sampled corals living at the most contrasted sites. These samples underwent next generation sequencing to reveal single-nucleotide-polymorphisms (SNPs) of which frequencies associated with heat stress gradients. As these SNPs might underpin an adaptive role, we characterized the functional roles of the genes located in their genomic neighborhood.

In each of the studied species, we found heat stress associated SNPs notably located in proximity of genes coding for well-established actors of the cellular responses against heat. Among these, we can mention proteins involved in DNA damage-repair, protein folding, oxidative stress homeostasis, inflammatory and apoptotic pathways. In some cases, the same putative molecular targets of heat stress adaptation recurred among species.

Together, these results underscore the relevance and the power of the seascape genomics approach for the discovery of adaptive traits that could allow corals to persist across wider thermal ranges.

## Introduction

One of the most dramatic consequences of climate change is the worldwide decline of coral reefs, which are the most biodiverse ecosystems in the marine environment (Hughes et al. 2017). Among the main drivers of this decline is coral bleaching, a stress response to anomalous heat waves that eventually causes the death of hard corals (Bellwood et al. 2004; Hughes et al. 2017). In the most severe episodes, coral bleaching provoked local living coral cover loss of up to 50% (Hughes et al. 2017; Hughes et al. 2018), with climate change projections expecting for bleaching conditions to be persistent worldwide by 2050 (Van Hooidonk et al. 2013).

Despite these catastrophic perspectives, a glimpse of hope is brought by coral reefs that show resistance after recurrent heat waves (Thompson and van Woesik 2009; Penin et al. 2013; Krueger et al. 2017; Dance 2019; Hughes et al. 2019). One of the mechanisms that might promote heat tolerance in corals is genetic adaptation (Sully et al. 2019). In recent years, there has been a growing body of literature investigating how coral thermal adaptation might alter the predictions of reef persistence, and how conservation policies could be modified accordingly (Logan et al. 2014; Van Oppen et al. 2015; Matz et al. 2018).

Given the crucial role adaptation will play in long-term reef persistence, there is an urgent need to characterize the adaptive potential of corals (Logan et al. 2014; Van Oppen et al. 2015). For instance, there are still open questions concerning the spatial and temporal scales at which local adaptation operates (Matz et al. 2018; Roche et al. 2018). Changes in adaptive potential have been observed along thermal gradients over hundreds of kilometres (*e.g*. (Thomas et al. 2017), but also at reefs with distinct thermal variations located only a few hundreds of meters apart (*e.g*. Bay and Palumbi 2014). Furthermore, different coral species are reported to show differential vulnerability against thermal stress, leading to the question of how different life-history traits (*e.g*. reproductive strategies) drive the pace of adaptation (Loya et al. 2001; Darling et al. 2012; Hughes et al. 2018).

There are also open questions concerning the molecular mechanisms that might be targeted by heat stress adaptation in corals (van Oppen and Lough 2009; Mydlarz et al. 2010; Palumbi et al. 2014). Some cellular responses to heat stress are now well characterized, such as DNA repair mechanisms, the activation of the protein folding machinery in the endoplasmic reticulum (ER) or the accumulation of reactive oxygen species (ROS, either endogenous or produced by the symbiont) that progressively elicits inflammatory and apoptotic responses (van Oppen and Lough 2009; Mydlarz et al. 2010; Maor-Landaw and Levy 2016; Oakley et al. 2017; Patel et al. 2018). However, little is known about which of the many molecular actors participating to these cascades could be hijacked by evolutionary processes to increment thermal tolerance.

Seascape genomics could contribute to filling these gaps. Seascape genomics is a budding field of population genomics that allows the study of local adaptation in wild populations (Riginos et al. 2016). This method combines the environmental characterization of the seascape with a genomic analysis of its population (Rellstab et al. 2015). The goal is to identify genetic variants that correlate with environmental gradients that might underpin an adaptive role (Rellstab et al. 2015). Seascape genomics could enhance the characterization of coral adaptive potential because: (1) it requires an extensive sampling strategy that allows for studying adaptation at different geographic scales, and against different types of environmental constraints simultaneously (*e.g*. mean temperatures, standard deviations, accumulated heat stress; Leempoel et al. 2017; Selmoni et al. 2020a); (2) its experimental protocol is less laborious in comparison to traditional approaches used for studying coral adaptation (*e.g*. aquarium experiments, transplantations), and therefore facilitates scaling-up to a multiple species analysis; (3) it is based on genomic data and thus reports candidate molecular targets of adaptation (Rellstab et al. 2015; Riginos et al. 2016). Moreover, recent work described how the results of seascape genomics studies on corals can be directly transposed to a conservation perspective and support reef prioritization (Selmoni et al. 2020b).

Here we applied the seascape genomics approach to study the adaptive potential against heat stress in three bleaching-prone coral species of New Caledonia, in the southwestern Pacific (Fig. 1). We first used publicly available satellite data to characterize the seascape conditions for over 1,000 km of the reef system. A sampling campaign was then organized to collect colonies at the 20 sites exposed to the most contrasted environmental conditions. The collected samples underwent a genotype-by-sequencing (DArT-seq) genomic characterization, followed by a seascape genomics analysis accounting for the confounding role of demographic structure. This allowed us to uncover single nucleotide polymorphisms (SNPs) associated with heat stress. We then performed the functional annotations of genes surrounding these SNPs and found molecular targets that notably recurred among species and that referred to well established heat stress responses in coral cells. Our study lays the foundations for the discovery of adaptive traits that could allow corals to persist across wider thermal ranges.

**Figure 1.**
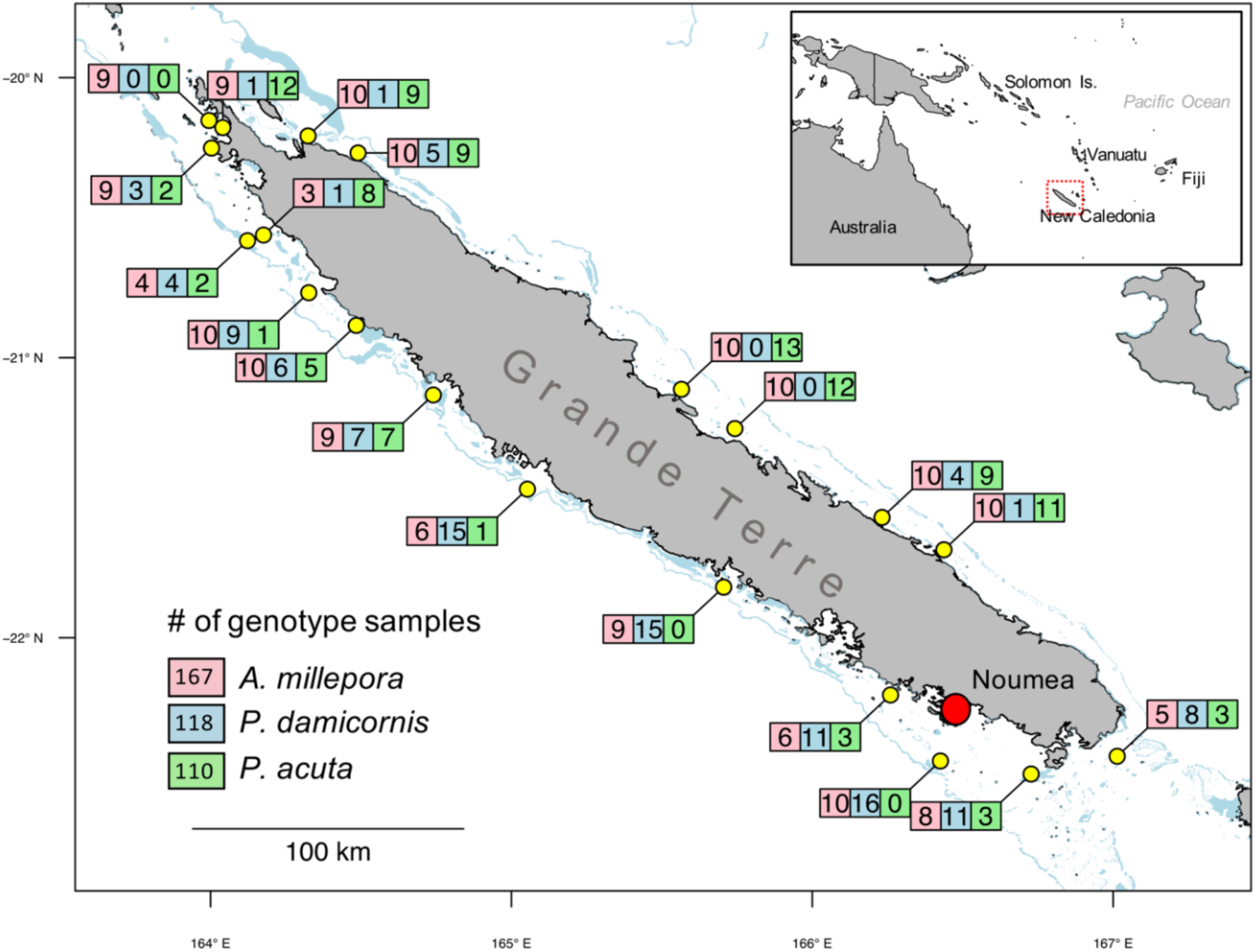
Study area and sampling sites. The 20 sampling sites around Grande Terre, the main island of New Caledonia (South Western Pacific), are shown in yellow. For every sampling site, the number of genotyped individuals per species (*Acropora millepora*: red, *Pocillopora damicornis*: blue, *Pocillopora acuta*: green) are given in the corresponding boxes.

## Results

Three coral species were sampled at 20 sites across the reef system of New Caledonia: *Acropora millepora* (Ehrenberg, 1834; n=360), *Pocillopora damicornis* (Linnaeus, 1758; (n=128) and *Pocillopora acuta* (Lamarck, 1816 ; n=150; Tab. S2). The DArT-seq analytical pipeline resulted in the genotyping of 188 samples by 57,374 bi-allelic single nucleotide polymorphisms (SNPs) for *A. millepora*, and 128 and 150 samples by 70,640 SNPs for *P. damicornis* and *P. acuta*, respectively (Tab. 1). After filtering for rare variants, missing values and clonality, we obtained a final genotype matrix of 167 individuals by 11,935 SNPs for *A. millepora*, of 118 individuals by 7,895 SNPs for *P. damicornis* and of 110 individuals by 8,343 SNPs for *P. acuta* (Tab. 1). The *A. millepora* genotyped samples distributed across all the 20 sampling sites (18 of which counted five samples or more), while genotyped samples of *P. damicornis* and *P. acuta* were distributed across 17 sites each (both with 10 sites counting five samples or more), with 15 sites where both species were found in sympatry (Fig. 1; Tab. S1).

**Table 1.** Workflow of the analysis. For each of the species of interest (*Acropora millepora, Pocillopora damicornis, Pocillopora acuta*), we report the number of individuals (ind.) and single nucleotide polymorphism (SNPs) obtained or retained after each the various step of the workflow.

|  | *A. millepora* | *Pocillopora* |  |
| --- | --- | --- | --- |
|  |  | *P. damicornis* | *P. acuta* |
| **Sampling** | 370 ind. | 360 ind. |  |
| **Microsatellite** | - | 128 ind. | 150 ind. |
| **DART-seq** | 188 ind. x 57,374 SNPs | 128 ind. x 70,640 SNPs | 150 ind. x 70,640 SNPs |
| **BLAST against reference** | 188 ind. x 47,529 SNPs | 127 ind. x 48,049 SNPs | 145 ind. x 48,049 SNPs |
| **Filtering (Missing values, MAF, MGF, LD, clonality)** | 167 ind. x 11,935 SNPs | 118 ind. x 7,895 SNPs | 110 ind. x 8,343 SNPs |

### Neutral genetic structure

We ran a principal component analysis (PCA) of the genotype matrix of each species to anticipate possible confounding role of neutral genomic variation on the adaptation study (Fig. 2, Fig. S1-S2). The Tracy-Widom test (*P*<0.05) revealed that the number of PCs underlying a non-random structure were seven for all the species, accounting for 8% of the total variance in *A. millepora*, 15% in both *P. damicornis* and *P. acuta* (Fig. 2). In *P. damicornis*, the spatial distribution of PC1 values appeared to be spatially structured following a north-south separation along the west coast of Grande Terre (Fig. S1B). In *P. acuta*, colonies in the north-west of Grande Terre displayed lower PC1 values, compared to those on the eastern coast (Fig. S1C). In *A. millepora*, no clear geographical patterns emerged as individuals with different values on PC1 were often located on the same reef. Finally, we analysed the presence of genomic widows clustering SNPs with high PC1-loadings, expected to be frequent in genetically isolated groups (cryptic species, hybrids). These genomics windows were rare, as we observed no more than three per species (Fig. S2).

**Figure 2.**
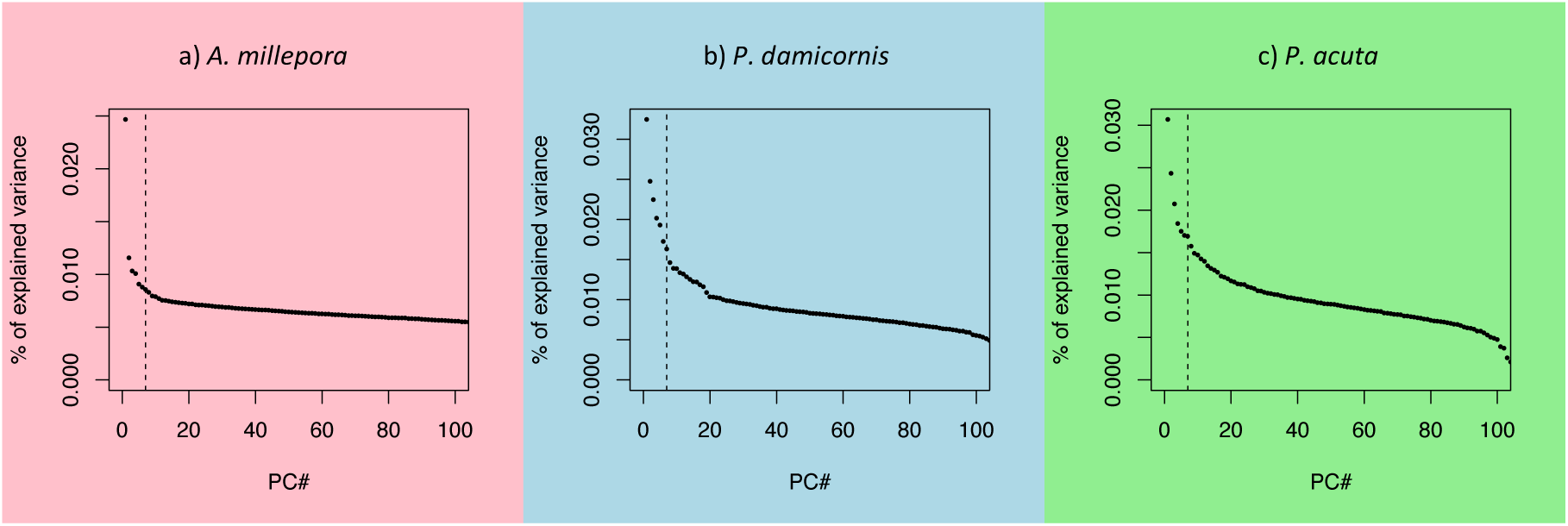
Principal component analysis (PCA) of the genotype matrices for three species studied, *Acropora millepora, Pocillopora damicornis* and *Pocillopora acuta*. The three graphs display the percentage of variance explained by the first 100 principal components (PC) of the genotype matrix for the three studied species. The vertical dotted lines represent the number of PCs deemed as underlying a non-random structure by the Tracy-Widom test (*P* < 0.05).

### Local adaptation

We investigated the presence of SNPs that were associated with 47 environmental gradients describing the seascape conditions in New Caledonia (Fig. 3, Fig. S3, Tab. S2). When a SNP was found associated with multiple environmental descriptors, only the most significant association was kept. In total, 120 significant (*q*<0.01) genotype-environment associations were found for *A. millepora*, 90 for *P. damicornis* and 100 for *P. acuta* (Tab. 2a; Tab. S3, S4). In all of the three species, we investigate the environmental descriptors that most frequently associated with significant SNPs were those related to sea surface temperature (SST; 63 in *A. millepora*, 47 in *P. damicornis*, 43 in *P. acuta*). Among these, we found that putative adaptive signals related to bleaching alert frequencies (73 genotype-environment associations) were more frequent than those relating to the standard deviation (42) and average temperatures (38; Tab. 2b).

**Table 2.**
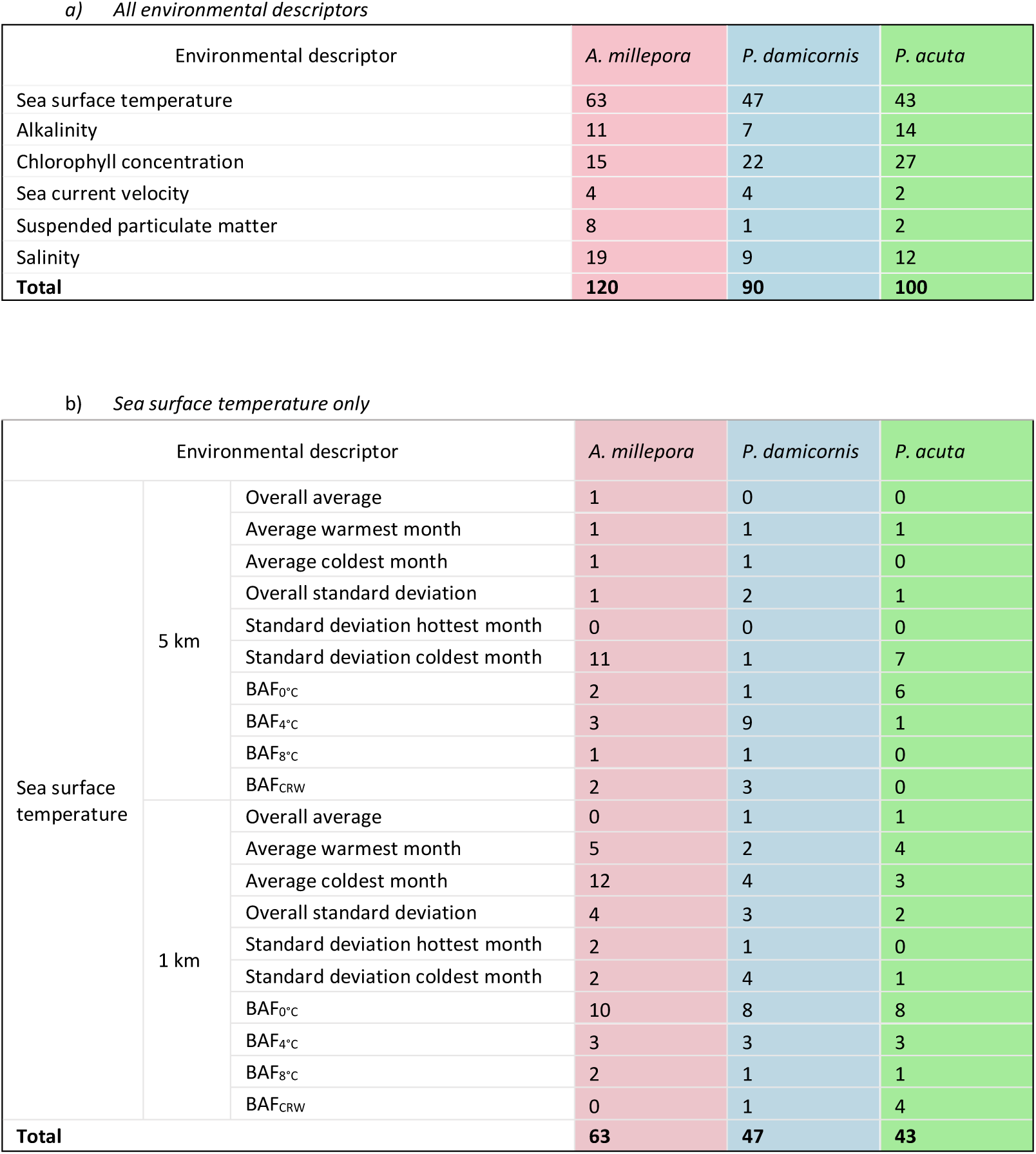
Significant genotype-environment associations in *Acropora millepora, Pocillopora damicornis* and *Pocillopora acuta*. Table a displays the number of SNPs significantly associated (*q*<0.01) with environmental descriptors; Table b, the complete list of environmental descriptors related to sea surface temperature (averages and standard deviations at two different spatial resolutions and indices of bleaching alert frequencies - BAF). Note that when a SNP was significantly associate to multiple environmental descriptors, only the best association was kept. The detailed list of the SNP-environment associations is available in the supplementary Table 4.

**Figure 3.**
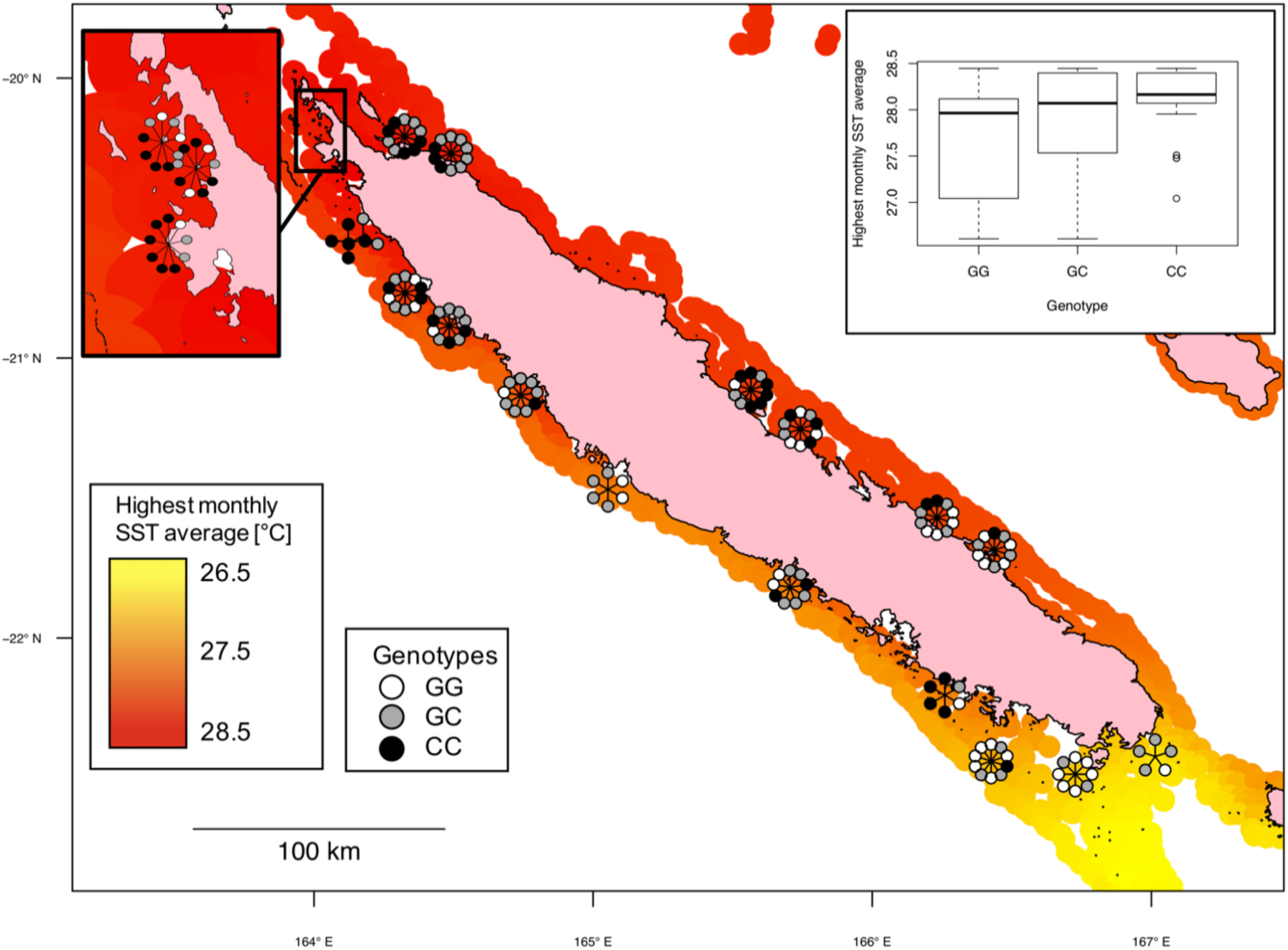
Example of significant genotype-environment association. The map displays the superposition between environmental gradient (here highest monthly SST average) and the distribution of an associated (q<0.01) SNP of *Acropora millepora*. Every circle corresponds to the SNP genotype for an individual colony. For illustrative reasons, genotypes are radially distributed around the sampling locations. The boxplot in the top-right corner shows how the environmental variable distributes within each genotype. The SNP represented here is located on the contig xpSc0000535 (position 118526) of *A. millepora* genome, and the closest annotated gene codes for ATP-dependent DNA helicase Q5.

When we focused on the environmental descriptors not relating to temperature, we observed that those describing chlorophyll concentration were those associated to more SNPs (64 across the three species), followed by salinity (40; Tab. 2a). In contrast, current velocity variables were the best environmental descriptors associated with fewer genetic variants (10 across the three species, Tab. 2a).

### Functional annotations of heat stress associated SNPs

Genes around SNPs that were associated with heat stress were annotated with gene ontology (GO) terms to investigate the molecular functions potentially altered by a genetic variant. GO terms were ranked for over-representation according to the Fisher exact test *P*-value (Tab. 3, S5). Among the top 50 ranks we found GO terms describing molecular functions such as “mismatched DNA binding”, “heat shock protein binding”, “chaperone binding”, “unfolded protein binding”, “cytoskeletal protein binding” and “actin binding” in *A. millepora* (Tab. 3a, S5a); “actin filament binding receptor activity”, “exonuclease activity”, “endonuclease activity” and “death effector domain binding” in *P. damicornis* (Tab. 3b, S5b); “NAD binding”, “nucleotide binding” and “mitogen-activated protein kinase binding” in *P. acuta* (Tab. 3c, S5c). Four terms (“signaling receptor binding”, “receptor regulator activity”, “organic cation transmembrane transporter activity”, “enzyme activator activity”) recurred among top ranked GO terms in at least two different species (Tab. 3). Each of the three species displayed top ranked GO terms referring to “oxidoreductase activity” acting on different molecules (Tab. 3).

**Table 3.** Functional annotations of heat stress associated SNPs. For each of the studied species, *Acropora millepora, Pocillopora damicornis* and *Pocillopora acuta*, the tables display the list of GO terms describing molecular functions that are overrepresented in the genomic neighborhoods (± 50 kb) of heat stress associated SNPs. For each GO term, the tables display the id (GO.ID), the term description (Term), the occurrence in the genome split in 100 kb windows (#Ann.), the observed (#Obs.) and expected (#Exp.) occurrence in the neighborhoods of heat stress associated SNPs and the *P*-value associated to the Fisher exact test comparing the expected and observed occurrences. For every species, the tables show a subset of the top 50 GO terms. The complete list of top ranked GO terms is available in the supplementary material (Tab. S5). GO terms in bold are those appearing in the top GO list of two different species.

| **a) *A. millepora*** |  |  |  |  |  |  |
| --- | --- | --- | --- | --- | --- | --- |
| Rank | GO.ID | Term | #Ann. | #Obs. | #Exp. | *P*-value |
| 3 | GO:0003779 | actin binding | 278 | 11 | 4.25 | <0.01 |
| 10 | GO:0016670 | oxidoreductase activity, acting on a sulfur group of donors, oxygen as acceptor | 10 | 2 | 0.15 | 0.01 |
| 12 | GO:0098772 | molecular function regulator | 724 | 19 | 11.07 | 0.01 |
| 15 | GO:0008047 | **enzyme activator activity** | 244 | 9 | 3.73 | 0.01 |
| 16 | GO:0030547 | receptor inhibitor activity | 11 | 2 | 0.17 | 0.01 |
| 25 | GO:0016701 | oxidoreductase activity, acting on single donors with incorporation of molecular oxygen | 38 | 3 | 0.58 | 0.02 |
| 26 | GO:0019209 | kinase activator activity | 39 | 3 | 0.6 | 0.02 |
| 32 | GO:0030983 | mismatched DNA binding | 16 | 2 | 0.24 | 0.02 |
| 33 | GO:0043028 | cysteine-type endopeptidase regulator activity involved in apoptotic process | 16 | 2 | 0.24 | 0.02 |
| 34 | GO:0008092 | cytoskeletal protein binding | 644 | 16 | 9.85 | 0.03 |
| 39 | GO:0015101 | **organic cation transmembrane transporter activity** | 19 | 2 | 0.29 | 0.03 |
| 41 | GO:0051082 | unfolded protein binding | 83 | 4 | 1.27 | 0.04 |
| 43 | GO:0060090 | molecular adaptor activity | 167 | 6 | 2.55 | 0.04 |
| 44 | GO:0016620 | oxidoreductase activity, acting on the aldehyde or oxo group of donors, NAD or NADP as acceptor | 22 | 2 | 0.34 | 0.04 |
| 47 | GO:0031072 | heat shock protein binding | 88 | 4 | 1.35 | 0.04 |
| 50 | GO:0030545 | **receptor regulator activity** | 131 | 5 | 2 | 0.05 |
| **b) *P. damicornis*** |  |  |  |  |  |  |
| Rank | GO.ID | Term | #Ann. | #Obs. | #Exp. | *P*-value |
| 2 | GO:0005506 | iron ion binding | 126 | 6 | 1.78 | 0.01 |
| 7 | GO:0016705 | oxidoreductase activity, acting on paired donors, with incorporation or reduction of molecular oxygen | 167 | 6 | 2.36 | 0.03 |
| 8 | GO:0051015 | actin filament binding | 127 | 5 | 1.79 | 0.03 |
| 11 | GO:0048487 | beta-tubulin binding | 24 | 2 | 0.34 | 0.04 |
| 15 | GO:0008047 | **enzyme activator activity** | 251 | 7 | 3.55 | 0.06 |
| 17 | GO:0004888 | transmembrane signaling receptor activity | 1045 | 20 | 14.76 | 0.06 |
| 19 | GO:0008528 | G protein-coupled peptide receptor activity | 324 | 8 | 4.58 | 0.08 |
| 20 | GO:0005102 | **signaling receptor binding** | 749 | 15 | 10.58 | 0.08 |
| 30 | GO:0005096 | GTPase activator activity | 139 | 4 | 1.96 | 0.13 |
| 40 | GO:0038023 | signaling receptor activity | 1142 | 20 | 16.13 | 0.14 |
| 42 | GO:0016796 | exonuclease activity, active with either ribo- or deoxyribonucleic acids and producing 5'-phosphomonoesters | 47 | 2 | 0.66 | 0.14 |
| 46 | GO:0016894 | endonuclease activity, active with either ribo- or deoxyribonucleic acids and producing 3'-phosphomonoesters | 11 | 1 | 0.16 | 0.15 |
| 47 | GO:0030169 | low-density lipoprotein particle binding | 11 | 1 | 0.16 | 0.15 |
| 49 | GO:0035877 | death effector domain binding | 11 | 1 | 0.16 | 0.15 |
| **c) *P. acuta*** |  |  |  |  |  |  |
| Rank | GO.ID | Term | #Ann. | #Obs. | #Exp. | *P*-value |
| 5 | GO:0022857 | transmembrane transporter activity | 702 | 18 | 10.17 | 0.01 |
| 9 | GO:0015368 | calcium:cation antiporter activity | 11 | 2 | 0.16 | 0.01 |
| 10 | GO:0005102 | **signaling receptor binding** | 749 | 18 | 10.85 | 0.01 |
| 21 | GO:0015101 | **organic cation transmembrane transporter activity** | 16 | 2 | 0.23 | 0.02 |
| 35 | GO:0000166 | nucleotide binding | 1297 | 25 | 18.79 | 0.03 |
| 41 | GO:0031435 | mitogen-activated protein kinase binding | 23 | 2 | 0.33 | 0.04 |
| 42 | GO:0030545 | **receptor regulator activity** | 137 | 5 | 1.98 | 0.05 |
| 45 | GO:0051287 | NAD binding | 57 | 3 | 0.83 | 0.05 |
| 47 | GO:0016651 | oxidoreductase activity, acting on NAD(P)H | 59 | 3 | 0.85 | 0.05 |
| 49 | GO:0090722 | receptor-receptor interaction | 26 | 2 | 0.38 | 0.05 |

## Discussion

### Different types of heat stress adaptation

In each of the studied species, we detected genotype-environment associations that might underpin local adaptation (Tab. 2a). Approximately half of these associations concerned descriptors of sea surface temperature (SST; Tab. 2a), partially because these are the most numerous types of descriptors employed in the analysis (20 out of 47; Tab. S2a).

As we focused on SST-related associations, we found that those involving bleaching alert frequencies (BAFs) were more frequent (73) than those related to temperature variations (42) and temperature averages (38; Tab. 2). Coral bleaching is a major threat for coral survival, and bleaching conditions emerge when SST variation exceeds seasonal averages (Liu et al. 2003; Hughes et al. 2017). BAFs descriptors account precisely for this selective constraint (SST variation over average), and this might explain why genotype-environment associations with BAFs were more frequent. Previous work on coral seascape genomics also reported a predominance of adaptive signals related to BAF (Selmoni et al. 2020b). Coral adaptation appeared to be also driven by SST averages (regardless of variations) or by SST variations (regardless of the averages; Tab. 2b). This kind of adaptation might relate to bleaching (*e.g*. being adapted to high thermal variability promotes bleaching resistance; Safaie et al. 2018), or to other types of heat stress responses (*e.g*. impaired injury recovery at elevated average SST; Bonesso et al. 2017).

### Candidate molecular targets for heat stress adaptation

Previous research reported that reefs exposed to high frequency of daily thermal variability showed reduced bleaching prevalence (Safaie et al. 2018). One of the reasons might be that corals at these sites manage to rapidly readjust their cellular homeostasis (Ruiz-Jones and Palumbi 2017). This view is supported by the numerous GO terms describing activity regulators (for instance “signaling receptor binding”, “receptor regulator activity”, “enzyme activator activity”, “molecular function regulator”; Tab. 3, S5) found surrounding heat stress associated SNPs. The fact that these terms are not heat stress specific, however, invites to a cautious interpretation.

In contrast, we also detected several genes coding for well-established molecular actors of corals thermal stress responses in the neighborhood of heat stress associated SNPs (Tab. 4, S4). In some rare cases, we found that SNPs fell directly in the coding sequence of genes, but more frequently the SNPs were located several kb distance from genes (Tab. 4). However, this does not exclude causative effects, as (1) the SNP detected could be physically linked to an adaptive SNP in the coding sequence; (2) the adaptive SNP could be located several kb from the target gene, as it is often the case (Brodie et al. 2016). Hereunder, we highlight the different molecular functions and the related proteins that were found as potential targets for thermal adaptation in corals.

**Table 4.**
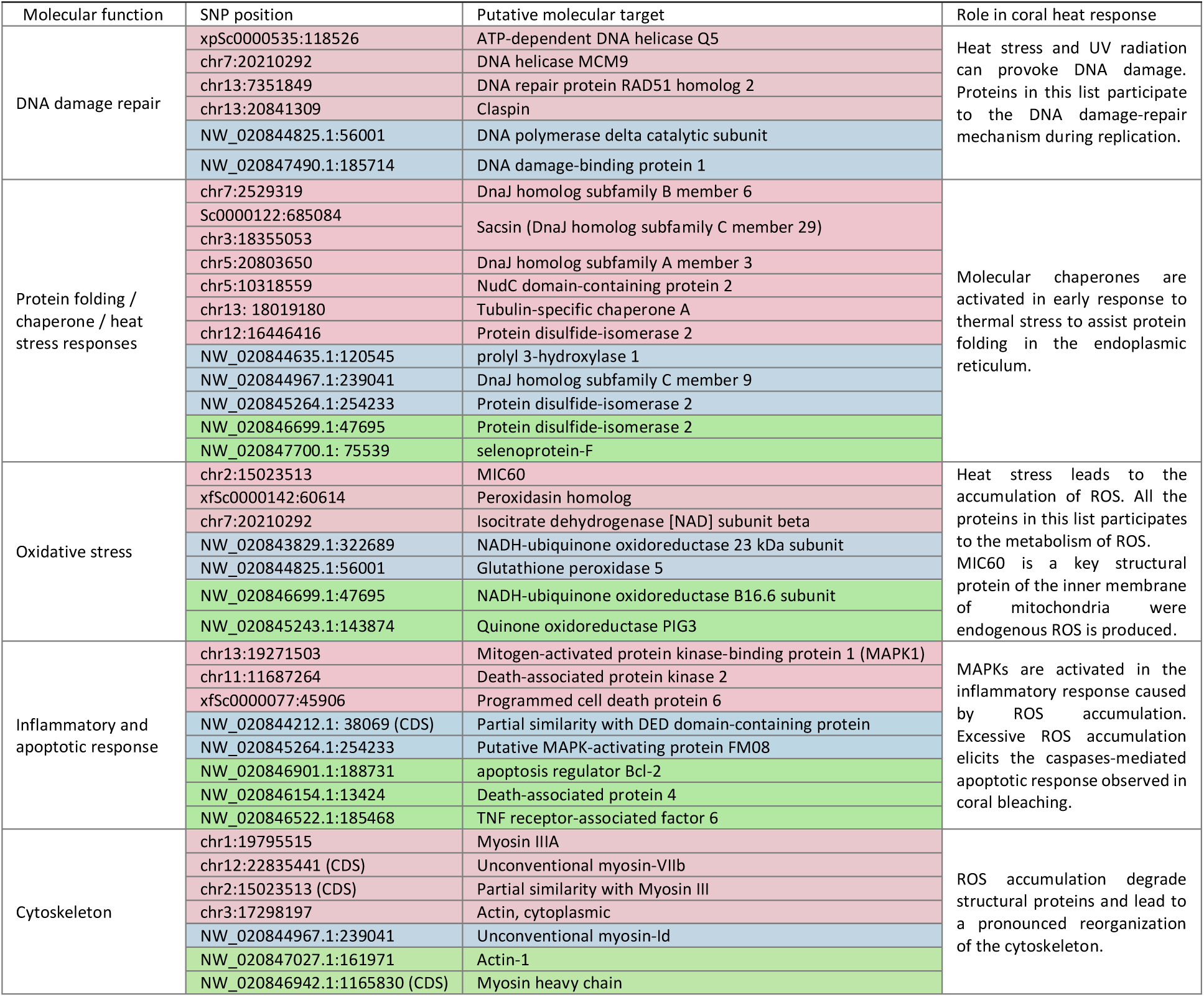
Candidate molecular targets for coral adaptation to heat stress. Annotations of genes surrounding (± 50 kb) heat stress associated SNPs were sorted by molecular function. For six specific types of molecular functions, the table displays the genes potentially involved in heat stress response (Putative molecular target) and the position of the corresponding SNP associated with heat stress (SNP position, in format contig/chromosome: base position). The CDS tag indicates SNP falling inside the coding sequence of the putative molecular target. The background colours correspond to the species where the candidate molecular target was found (pink: *Acropora millepora*, blue: *Pocillopora damicornis*, green: *Pocillopora acuta*). The role that each molecular function has in coral heat response is described in the last column.

### DNA repair

Heat stress impacts the integrity of nucleic acids and elicits mechanisms that promote DNA damage-repair and RNA stability (Henry et al. 1992; Sottile and Nadin 2018). A previous seascape genomics study on *Acropora digitifera* found five SNPs associated with heat stress to be proximal to genes coding for Helicase Q (Selmoni et al. 2020b). Here we found Helicase Q5 in the genomic neighbourhood of a SNP associated with heat stress in *A. millepora* (Tab. 4). Helicases Q are required for efficient DNA repair during the initiation of the replication fork (Sharma et al. 2006). Another family of helicases participating to this process are Helicases MCMs (Daniel et al. 2013) and one was found next to an heat stress associated SNP in *A. millepora* (Fig.3; Tab. 4).

In addition, we found proteins involved in DNA damage-repair that are known to be differentially expressed in corals under heat stress. For instance, claspin (Palumbi et al. 2014; Smits et al. 2019) and RAD51/54 homologs (Maor-Landaw and Levy 2016) were found here surrounding heat stress associated SNPs in *A. millepora*, while DNA damage-binding protein 1 (Li et al. 2006) and DNA polymerase delta catalytic subunit (Prindle and Loeb 2012) in *P. damicornis* (Tab. 4). Of note, GO terms describing molecular functions related to DNA repair (“mismatched DNA binding”, “exonuclase activity”, “endonuclease activity”, “nucleotide binding”) were found as over-represented in genes surrounding heat stress associated SNPs in the three species (Tab. 3).

### Protein folding

One of the main groups of gene annotations surrounding heat stress associated SNPs concerned molecular chaperones (Tab. 4). These are proteins that intervene in cellular responses to heat stress, where they assist the folding or unfolding of proteins in the endoplasmic reticulum notably (ER; Oakley et al. 2017). In corals, the role of these proteins in heat response, as well as their up-regulation under thermal stress, have been reported in several studies (Desalvo et al. 2008; Ishikawa et al. 2009; van Oppen and Lough 2009; Desalvo et al. 2010; Rosic et al. 2011; Maor-Landaw and Levy 2016; Oakley et al. 2017; Ruiz-Jones and Palumbi 2017). The annotation analysis for the related GO terms (*e.g*. “heat shock protein binding”, “unfolded protein binding”), revealed that this function was over-represented in genes close to the heat stress associated SNPs of *A. millepora* (Tab. 3a). In the three species, the genomic neighborhood of heat stress associated SNP contained several classes of chaperones: DnaJ homologs (four in *A. millepora*, one in *P. damicornis*), Tubulin-specific chaperone A and NudC domain-containing protein (*A. millepora*; Zheng et al. 2011), prolyl 3- hydroxylase 1 (*P. damicornis*; Ishikawa et al. 2009) and selenoprotein-F (*P. acuta;* Ren et al. 2018). Another important class of chaperones are “Protein disulfide-isomerase”, which catalyze the formation or breakage of disulfide bonds in proteins and produce reactive oxygen species (ROS) as byproducts (van Oppen and Lough 2009; Oakley et al. 2017). For example, the disulfide-isomerase gene expression has been shown to be upregulated in *Stylophora pistillata* under experimental heat stress (Maor-Landaw et al. 2014). Disulfide-isomerase 2 genes were found next to heat stress associated SNPs in each of the studied species (Tab. 4).

### Oxidative stress response

In parallel to the protein folding and recycling response, coral cells under heat stress accumulate ROS (Oakley et al. 2017; Nielsen et al. 2018). This accumulation can derive from the leakage of ROS from the damaged photosynthetic machinery of the endosymbiont, as well as from the endogenous production of the host mitochondria elicited under heat stress (Oakley et al. 2017; Nielsen et al. 2018). ROS accumulation causes oxidative stress, and the GO terms describing the inherent responses (“oxidoreductase activity”) were found as over-represented in genes next to heat stress associated SNPs in the three studied species (Tab. 3). For instance, we found genes coding for Peroxidasin homolog and Isocitrate dehydrogenase subunit beta next to heat stress associated SNPs in *A. millepora* (Tab. 4). Peroxidasin homolog is in the first line of defence against ROS accumulation, displays high rates of evolution in *A. millepora* and was found highly up-regulated under heat stress in *Monstastraea faveolata* (= *Orbicella faveolata*; Voolstra et al. 2009; Voolstra et al. 2011; Louis et al. 2017). Isocitrate dehydrogenase is one of the few sources of NADPH in the animal cell and was found upregulated in *E. pallida* under heat stress (Kültz 2005; Oakley et al. 2017). NADPH is an essential substrate to contrast ROS accumulation (Oakley et al. 2017; Patel et al. 2018) and another gene implicated in its metabolism, Quinone oxidoreductase PIG3, was found close to heat stress associated SNPs in *P. acuta* (Tab. 4; Zangar et al. 2004). In *P. damicornis*, we found the Glutathione peroxidase 5 gene, belonging to a family of well-characterized antioxidants contrasting ROS accumulation in corals (Nielsen et al. 2018).

In the host mitochondria, ROS production occurs in a series of redox reactions across the inner membrane (the electron transport chain; Lutz et al. 2015). One of the main components of this chain is the respiratory complex I (NADH-ubiquinone oxidoreductase), and we found genes coding for two of its subunits in the genomic neighborhood of heat stress associated SNPs in both *Pocillopora* species (Tab. 4). The mechanism leading to ROS leakage from host mitochondria into coral cell cytoplasm is poorly known (Dunn et al. 2012; Oakley et al. 2017; Nielsen et al. 2018). However, it is noteworthy to mention that in *A. millepora* the SNP most strongly associated with heat stress was close to the MIC60 gene (Tab. 4, S4a). MIC60 is a subunit of the MICOS complex, a key protein in the maintenance of the mitochondrial inner membrane architecture, through which ROS are produced, and the outer membrane, through which ROS diffuse into cytoplasm (Muñoz-Gómez et al. 2015; Zhao et al. 2019).

### Inflammatory response and apoptosis

The effects of ROS depends on the level of accumulation: medium levels elicit an inflammatory response, while excessive levels lead to cell apoptosis (Patel et al. 2018). Mitogen-activated protein kinases (MAPK) are key actors in the inflammatory response (Son et al. 2013; Courtial et al. 2017; Patel et al. 2018). In corals, MAPKs were shown to repress ROS accumulation in *S. pistillata* (Courtial et al. 2017) and were found in proximity of a SNP associated to heat stress in another seascape genomics study on Japanese *A. digitifera* (Selmoni et al. 2020b). Here we found a MAPK coding gene around heat stress associated SNPs in *A. millepora* (MAPK1), and genes coding for a MAPK activating protein *P. damicornis* (Putative MAPK-activating protein FM08) and *P. acuta* (TNF receptor-associated factor 6; Tab. 4; Mason et al. 2004).

ROS excess eventually results in cell apoptosis (Patel et al. 2018). In each of the three species, we found apoptosis-related genes near heat stress associated SNPs: Programmed cell death protein 6 and Death-associated protein kinase 2 (*A. millepora);* Death effector domain-containing protein (*P. damicornis*); and apoptosis regulator Bcl-2 and Death-associated protein 4 (*P. acuta*; Tab 4). These proteins participate in the caspase-mediated apoptotic cascade involved in the coral bleaching process (Ahmad et al. 1997; Dunn et al. 2007; Valmiki and Ramos 2009; Tchernov et al. 2011; Yuasa et al. 2015; Oakley et al. 2017).

### Cell structure

Heat stress has been shown to lead to cytoskeleton reorganization (Wilson et al. 2016). In Cnidaria, cytoskeletal proteins displayed changes in abundance under experimental heat stress in *Exaiptasia pallida* and *A. palmata* (Ricaurte et al. 2016; Oakley et al. 2017). Here we found several genes implicated in the cytoskeletal architecture in proximity of SNPs associated to heat stress: myosin III (twice), unconventional myosin VIIb and actin (*A. millepora*), unconventional myosin-Id (*P. damicornis*), Myosin heavy chain and actin-1 (*P. acuta*; Tab. 4). Moreover, the GO terms “actin binding” and “cytoskeletal protein binding” were over-represented in the set of genes neighbouring heat stress associated SNPs in *A. millepora*, and “actin filament binding” in *P. damicornis* (Tab. 3a-b). Of note, in three of the myosin genes the heat stress associate SNPs were found inside the coding sequence (Tab. 4, S4).

### Limitations and future directions

Seascape genomics studies are exploratory analyses that come with the drawback of being subjected to high false discovery rates (Rellstab et al. 2015; Riginos et al. 2016). This bias is stressed when the confounding role of neutral genetic variation is not accounted for (Selmoni et al. 2020a). The preliminary analysis of population structure, however, did not reveal any cryptic speciation nor isolated reefs among the studied populations (Fig. 2, S1-S2). Furthermore, we used a statistical method (LFMM) and a sampling design allowing to mitigate such confounding effects (Frichot et al. 2013; Selmoni et al. 2020a).

There are, nevertheless, some points that could have increased the statistical power of the analysis, as for instance the use of a larger sample size (over 200 individuals per species; (Selmoni et al. 2020a). In addition, the assignment of heat stress associated SNPs to candidate molecular targets for adaptation would have been further facilitated with a higher genome resolution in the sequencing strategy. Higher genome resolution would also allow to infer the structural modification that SNPs falling inside the coding sequence might cause. In the years to come, whole-genome-sequencing on corals is likely to become more affordable and can then be applied to the large sample sizes required for seascape genomics studies.

The next step in the characterization of corals’ adaptive potential is experimental validation. Our work found several genetic variants that might confer selective advantages against thermal stress (Tab. 4). For each of the studied species, we can now define multiple-loci genotypes of heat stress resistant colonies and test their fitness under experimental heat stress conducted in aquaria (Krueger et al. 2017). As a result, this analysis will allow to 1) further investigate the role of different heat stress associated genotypes and molecular pathways and 2) provide a concrete measure of the thermal ranges that these coral populations might sustain in the years to come. This information is of paramount importance, as it will allow to predict the reefs that are expected to already carry heat tolerant colonies and to define conservation strategies accordingly (Selmoni et al. 2020b). For instance, marine protected areas could be established to preserve reefs with higher adaptive potential against heat stress, where such reefs could provide the breeding stock to restore the damaged ones (Baums 2008; Van Oppen et al. 2015; van Oppen et al. 2017).

### Conclusions

In this study, seascape genomics allowed to uncover genetic variants potentially implicated in adaptive processes against different types of heat stress in three coral species of New Caledonia. These variants were located next to genes coding for molecular actors that participate in well-understood cellular reactions against thermal stress. In addition, the approach pointed out new candidate genes (*e.g*. Helicase Q) or processes (*e.g*. signalling receptor binding) that might be implied in such responses. Of note, some of these potential targets for adaptation recurred in the analyses of different species, supporting the robustness and the power of the seascape genomics. Future studies will focus on performing experimental assays to validate the implication of potentially adaptive genotypes and newly identified genes in the heat stress response and to measure the thermal ranges tolerated by the diverse adaptive genotypes.

## Material and methods

### Environmental data

The seascape genomics approach requires an exhaustive description of the environmental conditions in order to prevent the misleading effect of collinear gradients (Riginos et al. 2016; Leempoel et al. 2017). For this reason, the seascape characterization we used encompassed seven environmental variables: sea water temperature (SST), chlorophyll concentration, sea surface salinity, sea current velocity, suspended particulate matter, alkalinity and bleaching alert frequencies (BAF; Tab. S2). The environmental characterization was performed in the R environment (R Core Team 2016) using the *raster* package (Hijmans 2016) and following the method described in previous work on coral seascape genomics (Selmoni et al. 2020b) with some modifications outlined hereafter.

For the description of SST we used two different georeferenced datasets covering the extent of New Caledonia: (1) daily records of SST since 1981 at a spatial resolution of 5 km (SST_5km;_ EU Copernicus Marine Service 2017); (2) daily records of SST since 2002 at resolution of 1 km (SST_1km_; Group for High Resolution Sea Surface Temperature; Chao et al. 2009; Chin et al. 2017). The first dataset covers a wider temporal range, therefore providing a more reliable characterization of historical trends. The second dataset covers a smaller temporal window, but the higher geographic resolution allows to portray fine scale thermal patterns with a higher degree of confidence. Both datasets were used to compute, for each pixel of the study area, averages and standard deviations of the warmest month, the coldest month, and the entire observational period. Furthermore, both datasets were used to compute three indices of bleaching alert frequencies (BAF), representing the frequency of days (over the whole period of remote sensing) during which the bi-weekly accumulated heat stress (*i.e*. SST above the average maximum) exceeded 0°C (BAF_0°C_), 4°C (BAF_4°C_) and 8°C (BAF_8°C_) (Selmoni et al. 2020b). Similarly, we computed the frequency of the bleaching warning conditions as defined by the Coral Reef Watch (BAF_CRW_), corresponding to the accumulation of heat over the three previous months (Liu et al. 2003).

For the other datasets (chlorophyll concentration, sea surface salinity, sea current velocity and suspended particulate matter; EU Copernicus Marine Service 2017), the spatial resolution ranged between 4 and 9 km (Tab. S2). All the datasets covered a temporal extent of at least 20 years before 2018 (the year of sampling) and were processed to compute: (1) highest monthly average, (2) lowest monthly average and (3) overall average. For all the datasets captured at daily resolution (*i.e*. all except suspended particulate matter), we also computed the standard deviation associated with the three means. Seawater alkalinity was estimated by combining SST_5km_ and salinity in a polynomial equation as described by Lee and colleagues (Lee et al. 2006).

In total, 47 environmental descriptors (Tab. S2) were computed and assigned to the shapes of the reefs of New Caledonia (UNEP-WCMC et al. 2010), reported for a regular grid (∼3,000 cells of size: 2×2 km) using QGIS (QGIS development team 2009).

### Sampling

Twenty sampling sites were selected out of the ∼3,000 reef cells surrounding Grande Terre, the main island of New Caledonia (Fig. 1). Sampling sites were chosen following an approach that simultaneously maximized environmental contrasts and replicated them at distant sites (the “hybrid approach” described in Selmoni et al. 2020a). The method consists of applying Principal Component Analysis (PCA) and hierarchical clustering to the 47 environmental descriptors in order to separate the ∼3,000 reef cells into distinct environmental regions. Next, the algorithm selects the same number of sampling sites within each region in order to maximize physical distance between sites. Increasing environmental variation is expected to raise the sensitivity of seascape genomics analysis, while the replication of environmental gradients is expected to reduce false discovery rates (Selmoni et al. 2020a). Here the number of environmental clusters was five (Fig. S3) and we established at four sampling locations per cluster. When this was not possible (*e.g*. because of logistic constraints during the sampling campaign), additional sampling sites were added to the neighbouring clusters.

The sampling campaign was performed from February to May 2018 (under the permits N°609011-/2018/DEPART/JJC and N°783-2018/ARR/DENV) and targeted three flagship species of the Indo-Pacific: *Acropora millepora, Pocillopora damicornis sensu* Schmidt-Roach et al. (2013) [corresponding to PSH04 *sensu* P. Gélin et al. (2017)] and *Pocillopora acuta sensu* Schmidt-Roach et al. (2013) [corresponding to PSH05 *sensu* P. Gélin et al. (2017)]. Of note, *P. acuta* and *P. damicornis* belong to the complex of species formerly named *P. damicornis* (Schmidt-Roach et al. 2014; Johnston et al. 2017; Gélin, Fauvelot, et al. 2018). At every sampling site, we collected up to 20 samples of *A. millepora* and 20 of *Pocillopora* aff. *damicornis* (we did not discriminate between species while sampling as it can be difficult to distinguish them in the field). All the samples were collected in a 1 km area and at a depth ranging between 2-4 m. The centre of this area was used for georeferencing the sampling site. Before sampling, each colony was imaged underwater, then a portion of a branch was sampled with hammer and chisel. Each sample consisted of a 1-2 cm branch that was immediately transferred to 80% ethanol and stored at -20°C. DNA from the 730 samples (370 *A. millepora* and 360 *Pocillopora;* Tab. S1) were extracted using the DNeasy 96 Tissue kit (Qiagen) following manufacturer instructions.

### Pocillopora *species identification*

The 360 *Pocillopora* samples were identified molecularly *a posteriori* of sampling to be assigned to one species or the other (*P. damicornis or P. acuta*). Samples were thus genotyped using 13 microsatellite loci, as in Gélin et al. (2017; Online Resource 1). Then, colonies belonging to *P. damicornis* and to *P. acuta* were identified using assignment tests performed with Structure (v. 2.3.4 ;Pritchard et al. 2000), as in Gélin et al. (2018). Colonies assigned to *P. damicornis* or *acuta* with a probability of at least 0.70 were retained in the final dataset for this study. The *Pocillopora* sampling was composed of 148 *P. damicornis* (more precisely to SSH04b *sensu* Gélin et al. 2017), 159 *P. acuta* colonies (more precisely, a mix of SSH05a and SSH05b *sensu* Gélin, Pirog, et al. 2018) and 53 unassigned colonies (excluded from further analysis; Tab. S1).

### Acropora *species identification*

*Acropora* species are genetically and morphologically notoriously challenging in terms of identification and species boundaries detection. However, *A. millepora* can be recognised in the field based on its typical axial and radial corallite shape (Wallace 1999).Back from the field, *in situ* images of each specimen were examined to look for the species diagnostic morphological characters and initial identifications validated.

### Screening and SNP genotyping

All DNA samples from *A. millepora, P. damicornis* and *P. acuta* were sent to the Diversity Arrays Technology (Canberra, Australia) for quality check screening and genotype-by-sequencing using the DArT-sequencing method (DArT-seq; Kilian et al. 2012). The restriction enzymes used for library preparation for *A. millepora* and *Pocillopora* samples were PstI and HpaII. Prior to sequencing, all the DNA samples underwent a one-hour incubation with the digestion buffer, followed by a step of quality check for integrity, purity and concentration running 1 μL per sample on a 0.8% agarose gel. Samples from each site were then ranked based on their quality (degree of smearing on the agarose gel). We then selected the samples with the best scores from each site and defined a list of 188 *A. millepora*, 128 *P. damicornis* and 150 *P. acuta* samples that proceeded to the sequencing step in four and five lanes of an Illumina Hiseq2500, respectively. During each step of the workflow (DNA purification, library preparation and sequencing), *A. millepora* and *Pocillopora* samples were kept separated and randomly distributed across the respective batches (*e.g*. 96-well plates, sequencing lanes) to minimize the risks of technical bias. SNPs were called using the DArT-seq analytical pipeline (DArTsoft14).

### SNP filtering

The DArT-loci (*i.e*. the DNA sequences surrounding each SNP) initially underwent a sequence similarity search (BLASTn; v. 2.7.1; Madden and Coulouris 2008) against a reference genome to retain only those associated with the coral host. For *A. millepora*, the reference genome was the *A. millepora* chromosome-level assembly from Fuller and colleagues (v. 2; Fuller et al. 2019, unpublished data, available on arXiv), while for *P. damicornis* and *P. acuta* we used the only *Pocillopora* reference assembled to date (*P. damicornis sensu lato;* v. 1; Cunning et al. 2018). Only DArT-loci scoring an E-value below 10^−6^ were retained.

The processing of the SNPs data followed a pipeline from previous work on coral seascape genomics (Selmoni et al. 2020b). For each species’ dataset, we removed SNPs and individuals with high missing rates (> 50%) by using custom functions in the R environment. Next, we proceeded with imputation of missing genotypes using the *linkimpute* algorithm (based on k-nearest-neighbours imputation; Money et al. 2015) implemented in Tassel 5 (Bradbury et al. 2007) using the default settings. Afterwards, we repeated the filtering of SNPs and individuals for missing rates, but this time using a more stringent threshold (5%). We also applied a filter to exclude rare alleles (minor allele frequency < 5%) and highly frequent genotypes (major genotype frequency > 95%). SNPs were then filtered for linkage disequilibrium using the R package *SNPrelate* (function *snpsgdsLDpruning*, LDthreshold=0.3, v.1.16; Zheng et al. 2012). Finally, we applied a filter for clonality: when groups of colonies shared highly correlated genotypes (Pearson correlation > 0.9) only one colony pert group was kept.

### Neutral genetic structure analysis

Prior to the seascape genomics analysis, the neutral genetic structure of the studied populations was investigated by running a PCA on the genotype matrix of each species using the R *stats* package (*prcomp* function). Firstly, we visually inspected the percentage of variance of the genotype matrix explained by each principal component (eigenanalysis); in highly structured populations the first principal components (PCs) are expected to explain a larger proportion of the variance, when compared to the subsequent PCs (Johnstone 2001; Novembre et al. 2008). We ran a Tracy-Widom test as implemented in R package *AssocTests* (Wang et al. 2017) to determine the number of PCs underlying a non-random genetic structure (*P* < 0.05; Patterson et al. 2006).

We also visualized the spatial distribution of the main axis of variation (PC1), in order to investigate the presence of geographical structures (Novembre et al. 2008). Finally, we focused on the SNP-specific loadings on PC1 and their distributions across the genome. In fact, groups of genetically isolated individuals (*e.g*. hybrids, cryptic species) are expected to display genomic islands of low-recombination (*i.e*. groups of physically close SNPs contributing to high loading on the main axis of variation; Nosil et al. 2009; Li and Ralph 2019). We therefore visualized the distribution of average PC1-loadings by genomic windows of 50 and 100 kb. In these calculations, only genomic windows containing at least 5 SNPs were retained.

### Seascape genomics

The seascape genomics analyses were performed separately on the three species using the LFMM method implemented in the *LEA* R package (v. 2.4.0; Frichot et al. 2013; Frichot and François 2015). This method associates single environmental gradients and individual SNPs variations in mixed models, where the confounding effect of neutral genetic variation is accounted for through latent factors (Frichot et al. 2013).

Briefly, the first step of the LFMM pipeline is to estimate the number of latent factors (K; Frichot and François 2015). This parameter corresponds to the number of ancestral populations and can be estimated by using the *snmf* function of the LEA package. The method processes a genotype matrix to estimate individual admixture coefficients under different K’s, and then evaluates the quality of fit for each K via cross validation (Frichot and François 2015). We ran ten replicates of this analysis for all the studied species, and found that the optimal number of K (according to the lowest entropy criterion) ranged from 2-4 for *A. millepora*, 6-8 for *P. damicornis*, and 10-12 for *P. acuta* (Fig. S4).

We then proceeded to the genotype-environment association analysis with LFMM. Since this method does not accept missing genotypes, we first ran the *impute* function of the LEA package. For each species, this function inferred the missing genotypes out of the ancestral genotype frequencies previously calculated with the *snmf* function. Finally, we ran the association analysis between the SNPs of each species and the environmental condition descriptors. This was done by using the *lfmm* function, setting K to the ranges previously estimated for each species and running five replicates of each analysis. Since *lfmm* calculations can be computationally intensive, when two or more environmental descriptors were highly collinear (absolute value of Pearson correlation > 0.9), only one was used in the analysis.

LFMM returns *P*-values describing the statistical significance of every genotype-environment association under different values of K. For each association model related to the same environmental variable, *P*-values were corrected for multiple testing using the q-value method (R package *q-value*, v. 2.14, Storey 2003) and deemed significant if *q* < 0.01 under at least one level of K. When a SNP was significantly associated to multiple environmental variables, or with an environmental variable highly correlated with others (*i.e*. those excluded from the LFMM calculations), we defined a main environmental descriptor as the variable most strongly correlated (Pearson) with the SNP.

### Annotation analysis of heat stress associated SNPs

For each of the three studied species, we annotated the genomic neighbourhood of every SNP in order to characterize the possible functional implications of a genetic variant. Firstly, we uniformed the annotations of genes from different species in order to facilitate comparisons. We did this by retrieving the positions of genes in the two reference genomes (Cunning et al. 2018; Fuller et al. 2019) and the corresponding predicted protein sequences. These sequences were used to perform a similarity search (blastP; Madden and Coulouris 2008) against the Uniprot/swissprot database (*metazoa* entries, release 2020_01; Boeckmann et al. 2003). Each gene was annotated with the best significant hit (E-value < 0.01) and inherited protein name and gene ontology (GO) terms describing molecular functions when existing (Ashburner et al. 2000).

Afterwards, we focused on the annotation of genes surrounding SNPs associated with heat stress descriptors. Significant SNPs were deemed “heat stress associated” if they were best correlated with an environmental descriptor relating to average temperature, standard deviation of temperature, or bleaching alert frequency. We mapped every SNP associated with heat stress as the genes located within a ± 50 kb window. We selected this window size because genes associated with a SNP can be located hundreds of kilobases away (Visel et al. 2009; Brodie et al. 2016), with 50 kb being roughly the median contig size in the *P. damicornis* reference genome (Cunning et al. 2018).

We then computed the observed occurrence of each GO term among the genomic neighbourhoods of significant SNPs. As a comparison, we split the reference genome into 100 kb windows and computed the expected occurrence for each term. This procedure was performed separately for each of the three studied species, using the respective reference genome. The statistical analysis of differences between the expected and observed GO term occurrences (enrichment analysis) was performed using the Fisher exact test method as implemented in the R *topGO* package (v. 2.34; Alexa et al. 2006). As suggested in the *topGO* guidelines, we ranked GO terms according to the *P-value* of the Fisher test and discarded those occurring fewer than 10 times throughout the genome.

## Acknowledgments

We thank Gerard Mou-Tham, Joseph Baly and Miguel Clarque, for their support during the field campaign, Annie Guillaume for the comments and suggestions provided during the redaction of this paper and Andrew Baird for helpful discussion on field identifications.

This work was supported by the United Nations Environment Programme (UNEP) and International Coral Reef Initiative (ICRI) coral reefs small grants programme (grant number: SSFA/18/MCE/005). We also thank the Government of France and the Government of the Principality of Monaco who provided the funding for the small grants.

