## supplementary figures for "Seascape genomics reveals candidate molecular targets of heat stress adaptation in three coral species"

**Supplementary Figure 1. Spatial distribution of first principal component (PC1) of genotype matrix.** The PC1 values for the three studied species are represented on a scale from red (low value) to green (high value). Every point corresponds to an individual an its respective PC1 value. For illustrative reason, individuals are radially distributed around the sampling locations.

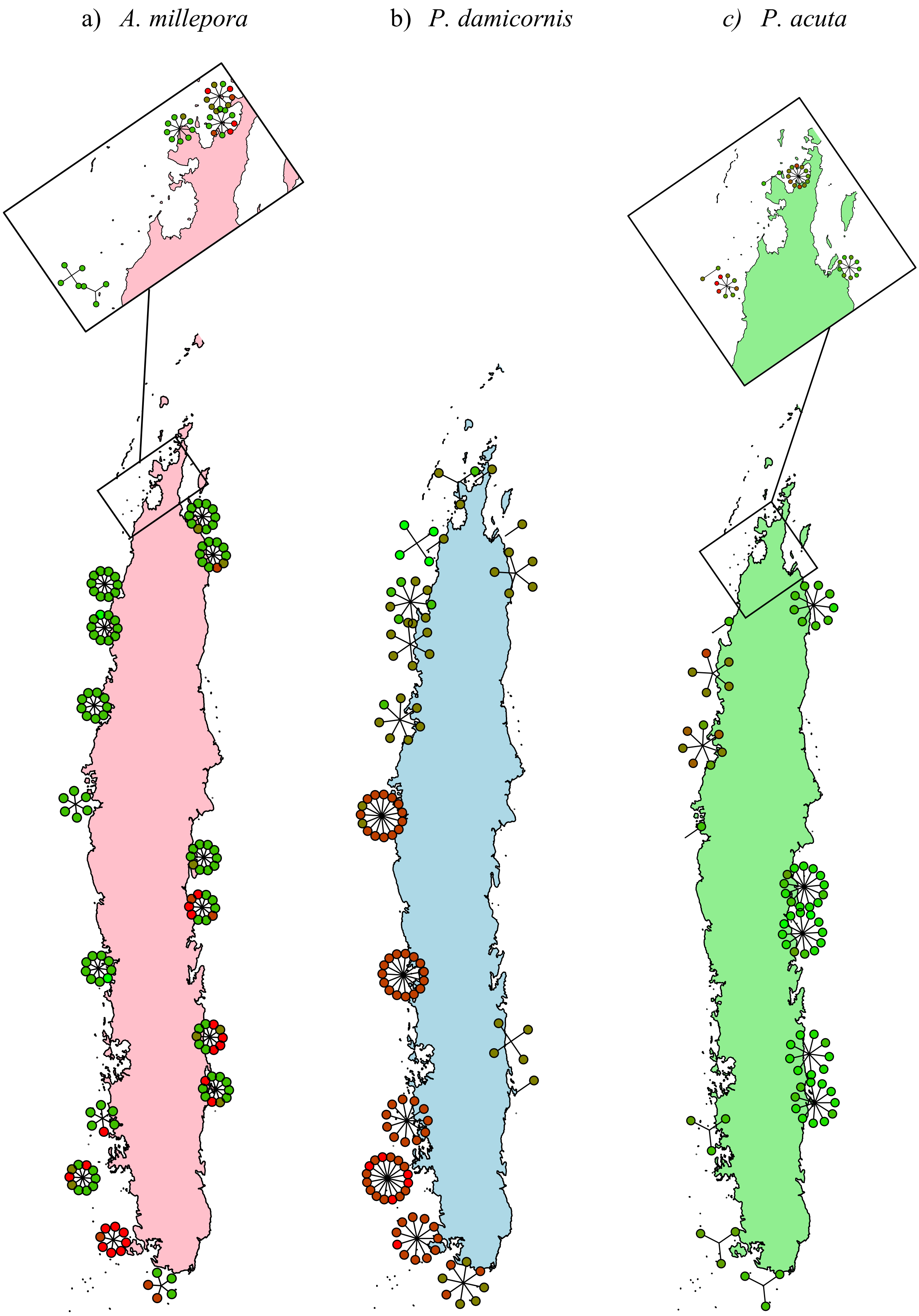

**Supplementary Figure 2. SNPs specific PC1-loadings averaged by genomic windows.** PC1-loadings for every SNP were averaged across genomics windows of 50 kbs (a, b, c) and 100 kbs (d, e, f). The histograms display the distributions of these averages for the three studied species.

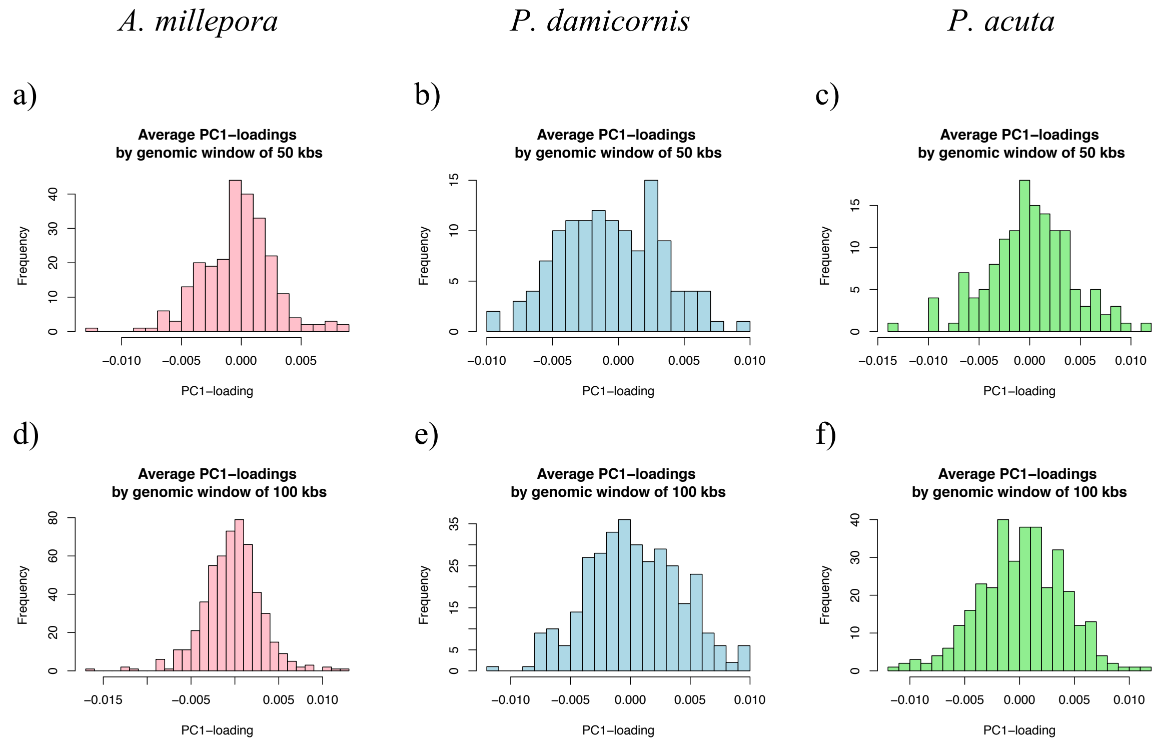

**Supplementary Figure 3. Environmental clusters of reefs of New Caledonia.** The reefs around Grande Terre are highlighted in five colors (blue, red, violet, orange and green) corresponding to the five clusters based on environmental variation. Where possible, we sampled corals at four sampling sites (yellow circles) per cluster.

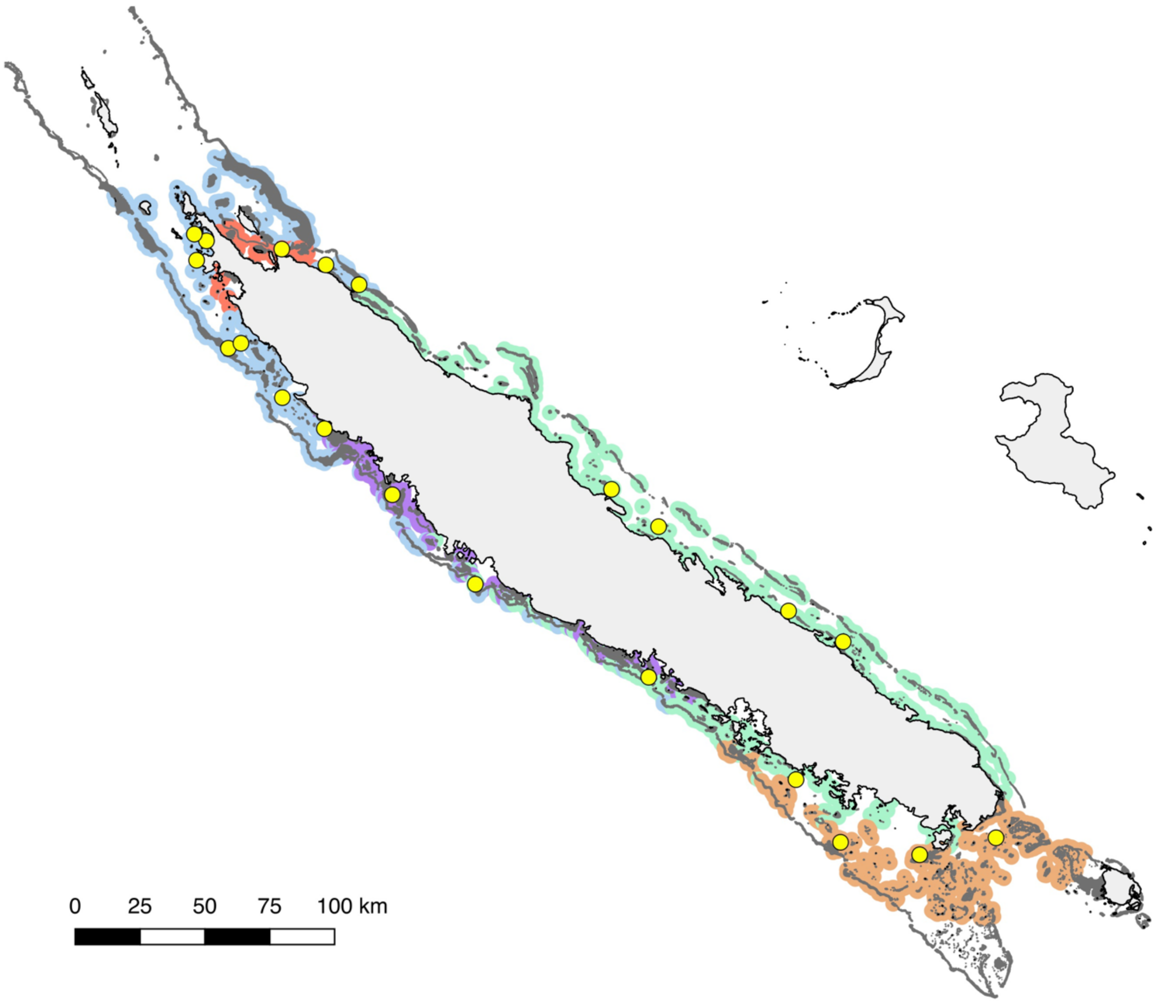

**Supplementary Figure 4. Cross-entropy comparison for the estimation of number of ancestral populations.** The graphs display the comparison of the quality of fit of admixture coefficients for different number of ancestral populations for the three species of interest. Lower cross-entropy criterion indicates a higher quality of fit.

a) *A. millepora*

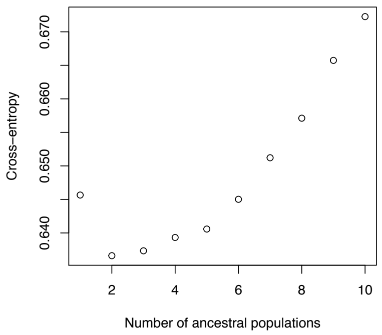


b) *P. damicornis*

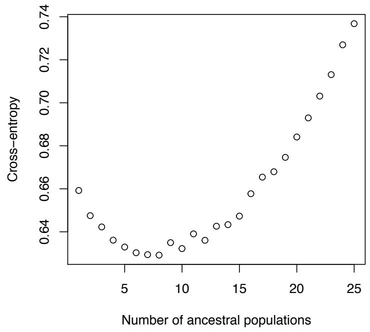


c) *P. acuta*

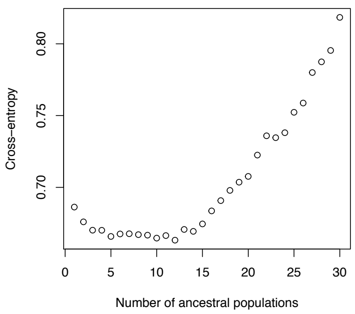
